# Operant self-stimulation of thalamic terminals in the dorsomedial striatum is constrained by metabotropic glutamate receptor 2

**DOI:** 10.1101/772525

**Authors:** Kari A. Johnson, Lucas Voyvodic, Yolanda Mateo, David M. Lovinger

## Abstract

Dorsal striatal manipulations including stimulation of dopamine release and activation of medium spiny neurons (MSNs) are sufficient to drive reinforcement-based learning. Glutamatergic innervation of the dorsal striatum by both the cortex and thalamus is a critical determinant of both MSN activity and local regulation of dopamine release. However, the relationship between glutamatergic inputs to the striatum and behavioral reinforcement is not well understood. We sought to evaluate the reinforcing properties of optogenetic stimulation of thalamostriatal terminals, which are associated with vesicular glutamate transporter 2 (Vglut2) expression, in the dorsomedial striatum (DMS), a region implicated in goal-directed operant behaviors. In mice expressing channelrhodopsin-2 (ChR2) under control of the Vglut2 promoter, brief optical stimulation of the DMS reinforces operant lever-pressing behavior. Mice also acquire operant self-stimulation of thalamic terminals in the DMS when ChR2 expression is virally targeted to the intralaminar thalamus. Because the presynaptic G protein-coupled receptor metabotropic glutamate receptor 2 (mGlu_2_) robustly inhibits glutamate and dopamine release induced by activation of thalamostriatal afferents, we examined the regulation of thalamostriatal self-stimulation by mGlu_2_. We find that administration of an mGlu_2/3_ agonist or an mGlu_2_-selective positive allosteric modulator reduces self-stimulation. In contrast, blockade of these receptors increases thalamostriatal self-stimulation, suggesting that endogenous activation of these receptors negatively modulates the reinforcing properties of thalamostriatal activity. These findings demonstrate that stimulation of thalamic terminals in the DMS is sufficient to reinforce a self-initiated action, and that thalamostriatal reinforcement is constrained by mGlu_2_ activation.

## INTRODUCTION

The dorsal striatum is a critical structure supporting reinforcement learning (Graybiel and Grafton, 2015). Classical intracranial self-stimulation (ICSS) studies in rats identified stimulation sites throughout the dorsal striatum that support operant responding (Phillips et al., 1976; Prado-Alcala and Wise, 1984). More recently, application of pathway-specific optogenetic techniques provided evidence that selective stimulation of nigrostriatal dopamine neurons is sufficient to support reinforcement learning (Ilango et al., 2014; Rossi et al., 2013). In addition to dopaminergic mechanisms, activation of striatal projection neurons (medium spiny neurons, MSNs) in either the medial (DMS) or lateral (DLS) subregion of the dorsal striatum reinforces actions (Kravitz et al., 2012; Lalive et al., 2018; Vicente et al., 2016). Because MSN activity critically depends on glutamatergic inputs, it is necessary to consider how glutamatergic projections to the striatum support reinforcement learning. Currently, whether activation of various glutamatergic inputs to the dorsal striatum is sufficient to drive reinforcement remains unclear.

MSNs receive major glutamatergic inputs from a variety of cortical and thalamic regions (Hintiryan et al., 2016; Hunnicutt et al., 2016; Smith et al., 2014). Whereas corticostriatal involvement in reinforcement learning has been studied extensively (Balleine and O’Doherty, 2010), relatively few studies have examined the contributions of thalamic projections to the striatum (Bradfield et al., 2013). Projections to the striatum from rostral intralaminar thalamic nuclei and from the parafascicular nucleus of the thalamus provide excitatory inputs to MSNs (Doig et al., 2010; Ellender et al., 2013; Huerta-Ocampo et al., 2014; Lacey et al., 2007). In addition, thalamic inputs to striatal cholinergic interneurons (CINs) locally control dopamine release via acetylcholine-mediated activation of nicotinic receptors on dopamine neuron terminals (Cover et al., 2019; Threlfell et al., 2012). The combined actions of thalamostriatal pathways on MSNs and dopamine release in the striatum suggest that activation of these projections could support behavioral reinforcement. Consistent with this idea, electrical stimulation of intralaminar thalamic nuclei that provide major projections to the dorsal striatum (including centromedial, centrolateral, and parafascicular nuclei) supports ICSS (Clavier and Gerfen, 1982). Additional evidence comes from a recent report that optogenetic stimulation of rostral intralaminar thalamic terminals in the dorsal striatum is sufficient to maintain operant responding in mice that were previously trained to respond for a different outcome (Cover et al., 2019). However, it remains unknown whether stimulation of thalamostriatal terminals in the DMS is sufficient to reinforce a novel, self-initiated action.

Presynaptic G protein-coupled receptors (GPCRs) expressed on glutamatergic terminals in the striatum regulate synaptic transmission and exert substantial influence over behavioral reinforcement (Atwood et al., 2014; Johnson and Lovinger, 2016). Agonists of the G_i/o_-coupled group II metabotropic glutamate (mGlu) receptors (mGlu_2_ and mGlu_3_) produce particularly robust presynaptic inhibition of glutamate release that is exclusively mediated by mGlu_2_ (Johnson et al., 2017; Kahn et al., 2001; Kupferschmidt and Lovinger, 2015; Lovinger and McCool, 1995; Martella et al., 2009). In addition, group II mGlu receptor activation reduces both basal and drug-evoked extracellular dopamine levels in the striatum (Bauzo et al., 2009; D’Souza et al., 2011; Hu et al., 1999; Kim et al., 2005; Pehrson and Moghaddam, 2010), although the mechanisms by which mGlu_2_ (and possibly mGlu_3_) constrain dopamine release have not been fully elucidated. Our recent work demonstrated that in addition to producing strong inhibition of both corticostriatal and thalamostriatal glutamatergic transmission in MSNs, selective activation of mGlu_2_ also dampens thalamostriatal glutamatergic transmission in CINs and reduces acetylcholine-mediated dopamine release evoked by thalamostriatal stimulation (Johnson et al., 2017). Thus, mGlu_2_ is poised to influence behavioral reinforcement associated with thalamically-driven glutamatergic and dopaminergic transmission in the dorsal striatum.

In this study, we evaluated operant responding for optical self-stimulation of thalamic terminals in the DMS. We report that mice readily press a lever for optical stimulation of thalamic terminals in the DMS without prior training for an alternative outcome or manipulation of motivational state. Consistent with our previous report that mGlu_2_ activation reduces glutamate and dopamine release driven by optical stimulation of thalamostriatal terminals (Johnson et al., 2017), group II mGlu receptor activation reduces thalamostriatal self-stimulation, whereas mGlu_2/3_ receptor blockade increases self-stimulation. These results support the notion that local manipulations of the DMS that produce MSN excitation and dopamine release are sufficient to drive reinforcement learning.

## RESULTS

### Optical stimulation of ChR2-expressing terminals in the DMS of Vglut2-Cre^+/−^;Ai32^+/−^ mice reinforces operant responding

Glutamatergic thalamic neurons that project to the dorsal striatum express the vesicular glutamate transporter Vglut2 (Fremeau et al., 2004). To examine the reinforcing properties of stimulation of Vglut2^+^ terminals in the DMS, we expressed ChR2 in Vglut2^+^ projections to the striatum by crossing Vglut2-IRES-Cre^+/−^ mice with Ai32^+/+^ mice (**Fig. 1a**). We implanted bilateral optical fibers in the DMS of Vglut2-Cre^+/−^;Ai32^+/−^ mice (**Fig. 1b, S1a-b**), then trained the mice to perform a self-paced self-stimulation task in an operant chamber with one lever. Lever presses were continuously reinforced with an optical stimulation train (20 pulses of 473 nm light delivered at 20 Hz). Across six training sessions, optical stimulation robustly reinforced pressing behavior in Vglut2-Cre^+/−^;Ai32^+/−^ mice, but not in Vglut2Cre^−/−^;Ai32^+/−^ mice (two-way RM ANOVA, significant session × genotype interaction, F_(5,45)_ = 8.105, p < 0.0001) (**Fig. 1c**). In Vglut2-Cre^+/−^;Ai32^+/−^ mice, lever presses per session were significantly higher in sessions 2-6 compared with session 1 (for each session 2-6 vs. session 1, p < 0.001, Dunnett’s multiple comparisons test). In contrast, we did not observe a significant escalation of lever pressing across training sessions in Vglut2Cre^−/−^;Ai32^+/−^ mice. Qualitatively, we observed that mice pressed the lever in clusters interspersed with periods in which the mice were not engaging with the lever (**Fig. 1d**).

**Figure 1.**
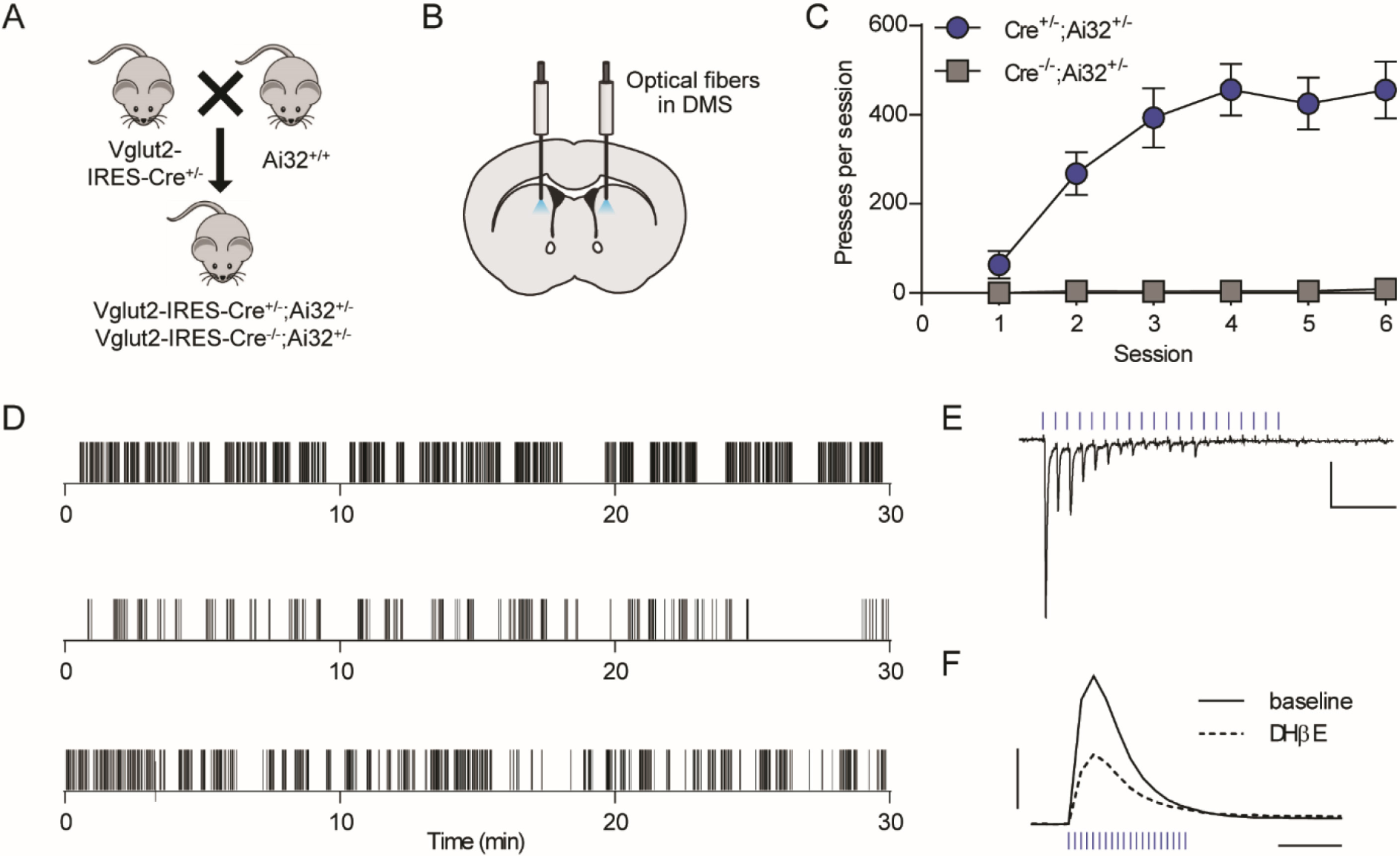
Optical stimulation of Vglut2^+^ terminals in the DMS reinforces operant lever pressing. (a) Breeding scheme. (b) Schematic of optical fiber placement in the DMS. (c) Acquisition of lever pressing for 20 pulses of 20-Hz optical stimulation in Vglut2-Cre^+/−^;Ai32^+/−^ (n = 7) and Vglut2-Cre^−/−^;Ai32^+/−^ (n = 4) mice. Data represent mean ± SEM. (d) Examples of lever presses from 3 representative Vglut2-Cre^+/−^;Ai32^+/−^ mice during acquisition session 6. Tick marks represent individual lever presses. (e) Representative whole-cell recording of EPSCs in a DMS medium spiny neuron in response to 20 pulses of 20-Hz optical stimulation. Scale bars: 100 pA, 0.25 s. (f) Representative FSCV traces of dopamine release measured in the DMS in response to 20 pulses of 20-Hz optical stimulation, before and after bath application of DHβE (1 µM). Scale bars: 0.5 µM dopamine, 0.5 s.

We examined responses to 20-Hz optical stimulation in brain slices to determine what synaptic mechanisms were activated by this protocol. The stimulation train produced excitatory postsynaptic currents (EPSCs) in MSNs recorded from slices of DMS, albeit with some failures in response to later light pulses in each train (**Fig. 1e**). This 20-Hz stimulation also evoked dopamine release in the DMS as measured by fast-scan cyclic voltammetry (FSCV) (**Fig. 1f**). Light-evoked dopamine release was partially blocked by the nicotinic receptor antagonist DHβE (1 µM) (**Fig. 1f, S1c**), suggesting that about half of light-evoked dopamine release in this model was mediated by CIN-derived acetylcholine actions on dopamine terminals, while the other half was due to either direct activation of dopaminergic afferents or another indirect effect.

### Pharmacological manipulation of mGlu_2_ modulates lever pressing for stimulation of Vglut2^+^ terminals in the DMS

The presynaptic GPCR mGlu_2_ robustly modulates glutamatergic and dopaminergic transmission in the dorsal striatum (Johnson et al., 2017). Thus, we predicted that pharmacological manipulation of mGlu_2_ could alter the reinforcing properties of Vglut2^+^ terminal stimulation in the DMS. Injection of the mGlu_2/3_-preferring antagonist LY341495 prior to the self-stimulation session did not significantly alter the number of active lever presses per session (119.3 ± 14.3% of vehicle, p = 0.25, paired t-test) (**Fig. 2a, d-e**). We found a dose-dependent decrease in pressing following injection of the mGlu_2/3_ agonist LY379268; 1 mg/kg did not significantly reduce pressing (72.1 ± 13.8% of vehicle, p = 0.10) (**Fig. 2b**,**d**), whereas we saw a substantial decrease in pressing following a 3 mg/kg dose (63.3 ± 6.8% of vehicle, p = 0.013) (**Fig. 2c-d, f**). To further understand the effects of mGlu_2/3_ activation on patterns of pressing, we quantified the number of clusters of pressing per session, the length of breaks from engaging with the lever, and within-cluster parameters including number of presses, duration, and press rate. Reduced lever pressing after LY379268 (3 mg/kg) administration was primarily driven by an increase in the length of breaks between clusters of pressing (**Fig. 2k**) without accompanying changes of within-cluster parameters or the number of clusters per session (**Fig. 2g-j**). In DMS brain slices prepared from Vglut2-Cre^+/−^;Ai32^+/−^ mice, optically-evoked dopamine release was reduced by LY379268 (100 nM); this effect was occluded by prior application of DHβE (1 µM), suggesting that LY379268 exclusively attenuates dopamine release driven by glutamatergic inputs to CINs (**Fig. S1d**).

**Figure 2.**
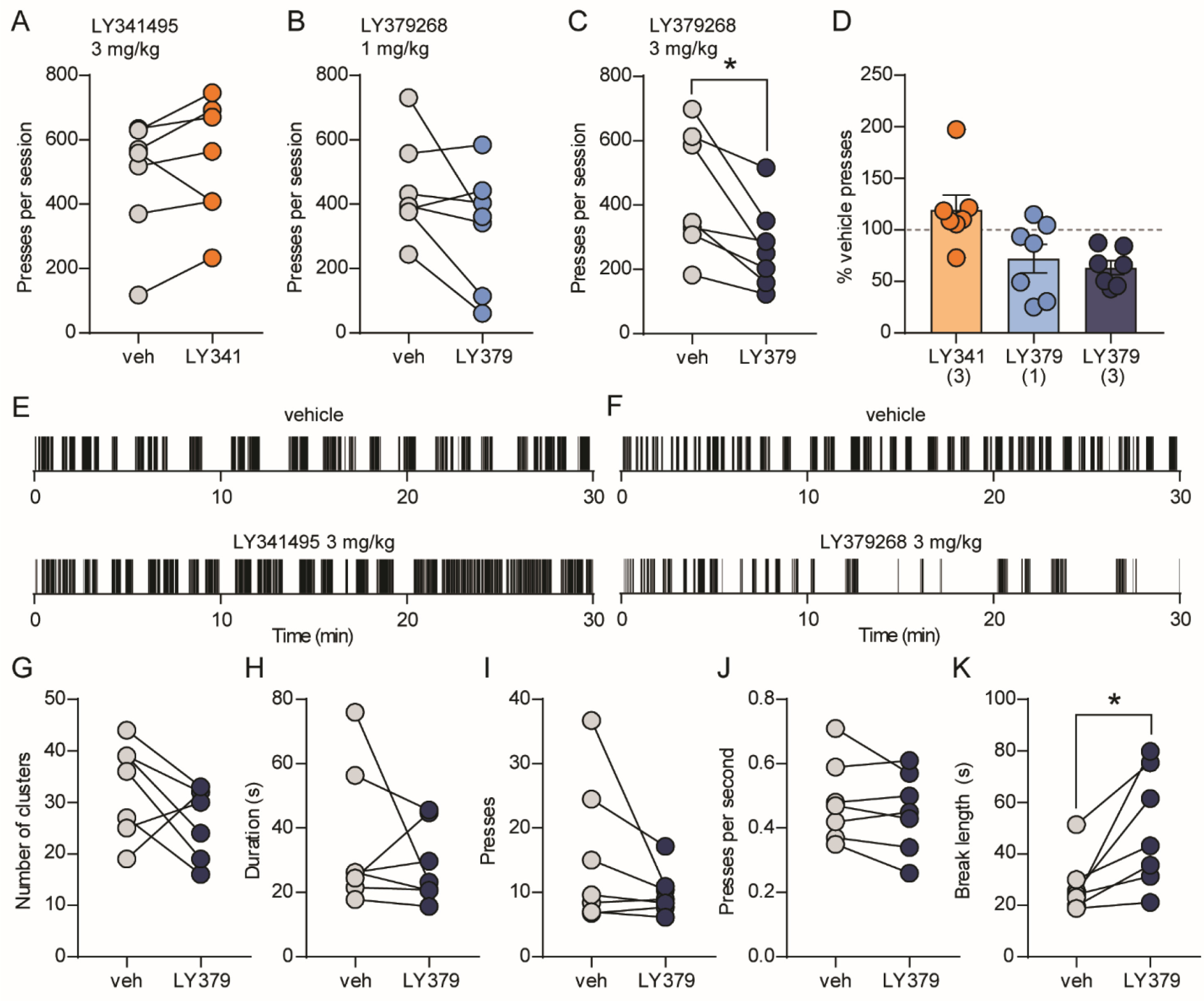
mGlu_2/3_ activation reduces operant responding for optical stimulation of Vglut2^+^ terminals in the DMS. (a-c) Within-subject comparisons of lever presses per session in Vglut2-Cre^+/−^;Ai32^+/−^ mice (n = 7) after injection of vehicle or 3 mg/kg LY341495 (a), 1 mg/kg LY379268 (b), and 3 mg/kg LY379268 (c). (d) Average lever presses per session (normalized to vehicle presses for each mouse) for each drug treatment. Bars represent mean ± SEM, and individual data points are overlaid. (e-f) Examples of lever presses from a representative mouse during vehicle or drug sessions. Tick marks represent individual lever presses. (g-k) Within-subject comparisons of patterns of pressing during vehicle or LY379268 sessions. Parameters analyzed were clusters of pressing per session (g), mean duration of clusters (h), mean number of presses per cluster (i), mean within-cluster press rate (j), and the mean length of breaks between clusters of pressing (k). *p < 0.05, paired t-test.

We implanted an additional cohort of Vglut2-Cre^+/−^;Ai32^+/−^ mice with optical fibers in the DMS (**Fig. S2a**) and trained the mice to distinguish between an active lever and an inactive lever to receive a stimulation train consisting of 10 pulses at 10 Hz, which more reliably produced EPSCs in MSNs (**Fig. S2b**) in addition to evoking dopamine release (**Figs. S1c, S2c**). Over the course of six training sessions, mice escalated pressing of the active lever but not of the inactive lever (**Fig. 3a**), and active lever presses per session in this paradigm were similar to numbers observed in mice trained with a single lever and receiving 20-Hz stimulation (**Fig. 1e**). Two-way RM ANOVA revealed a significant session × lever interaction (F_(5,25)_ = 18.76, p < 0.0001). For the active lever, total presses were higher during each of sessions 2-6 than during session 1 (p < 0.0001 sessions 2-6 vs. session 1, Dunnett’s multiple comparisons test). Inactive lever presses were not significantly different during sessions 2-6 compared with session 1. To determine the specific contribution of mGlu_2_ to modulation of responding for stimulation of ChR2-expressing terminals in the DMS, we compared the number of active lever presses following injection of the mGlu_2_-selective positive allosteric modulator BINA (15 mg/kg, i.p.) or vehicle. Like LY379268, BINA robustly decreased responding (46.6 ± 12.2% of presses during vehicle session, p = 0.0062, paired t-test) (**Fig. 3b**), confirming a specific role for mGlu_2_ in modulating reinforcement. No change in inactive lever pressing was observed.

**Figure 3.**
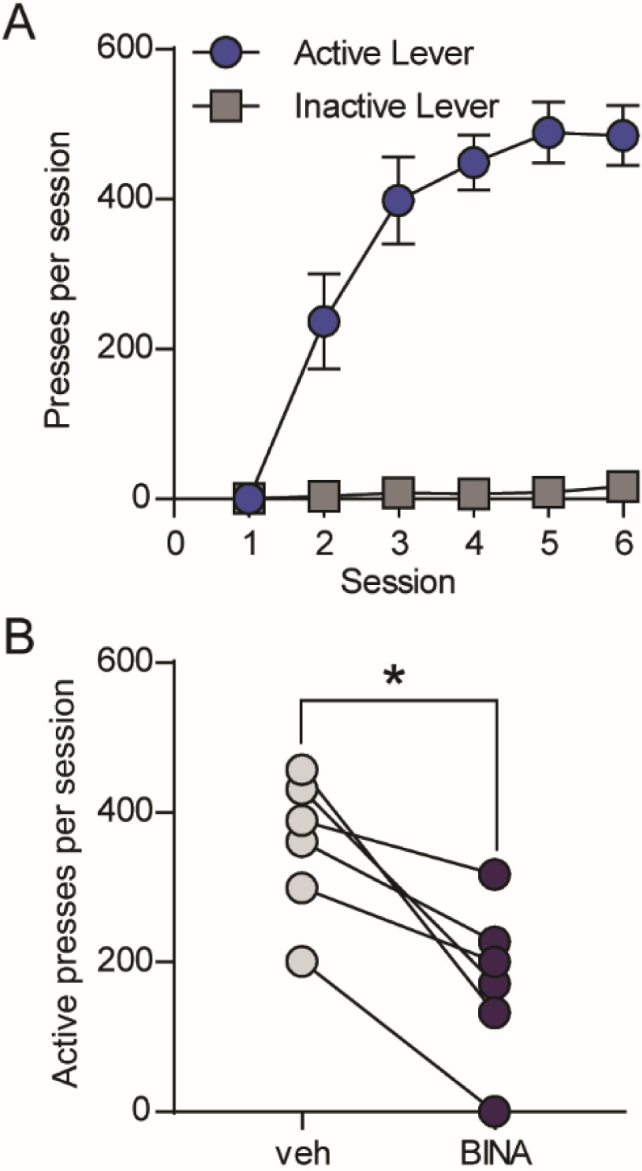
The mGlu_2_-selective positive allosteric modulator BINA reduces operant responding for optical stimulation of Vglut2^+^ terminals in the DMS. (a) Active and inactive lever pressing for 10 pulses of 10-Hz optical stimulation in Vglut2-Cre^+/−^;Ai32^+/−^ (n = 6). Data represent mean ± SEM. (b) Within-subject comparison of active lever presses per session after injection of vehicle or 15 mg/kg BINA (n = 6). *p < 0.05, paired t-test.

Previous studies in rats have shown that activation of mGlu_2/3_ modestly reduces operant responding for natural reinforcers such as palatable foods (Justinova et al., 2016; Kufahl et al., 2011). To assess the specificity of pharmacological manipulation of mGlu_2/3_, we trained food-restricted C57BL/6J mice to press a lever for delivery of a food pellet (**Fig. S3a-b**). In contrast to the lack of effect on Vglut2^+^ terminal self-stimulation, the mGlu_2/3_ antagonist LY341495 robustly reduced lever pressing for a food reinforcer to 49.7 ± 5.0% of vehicle (p = 0.0002, paired t-test) (**Fig. S3c**,**f**). Similar to previous reports in rats, the agonist LY379268 (1-3 mg/kg) produced modest decreases in pressing for a food reinforcer (1 mg/kg: 87.5 ± 3.3%, p = 0.0197; 3 mg/kg: 87.1 ± 5.1% of vehicle, p = 0.0371) (**Fig. S3d-f**).

### Optical stimulation of thalamic terminals in the DMS is sufficient to reinforce operant behavior

Although Vglut2^+^ terminals in the dorsal striatum are typically attributed to thalamic afferents (Fremeau et al., 2004; Smith et al., 2014), and little co-localization with dopamine neuron markers has been observed in the adult SNc (Poulin et al., 2018), our finding that optically-evoked dopamine release in the DMS of Vglut2-Cre^+/−^;Ai32^+/−^ mice is only partially sensitive to nicotinic receptor blockade suggests that ChR2 can stimulate dopamine release by an alternative mechanism in these mice, likely a direct effect on dopaminergic terminals. This result is consistent with reports of broader Vglut2 expression in dopamine neurons during development (Poulin et al., 2018; Trudeau et al., 2014). In addition, Vglut2^+^ inputs to CINs arising in the pedunculopontine nucleus were recently reported (Assous et al., 2019) and could also contribute to CIN-dependent dopamine release. To more specifically evaluate the reinforcing properties of thalamostriatal terminal stimulation, we virally expressed ChR2 bilaterally in the intralaminar nuclei of the thalamus and implanted bilteral optical fibers in the DMS of C57BL/6J mice (**Figs. 4a, S4b-c**), then trained mice in a self-paced self-stimulation task with an active and inactive lever available (**Fig. S4a**). Depression of the active lever resulted in optical stimulation consisting of 10 pulses of blue light delivered at 10 Hz. Over the course of nine training sessions, mice escalated pressing of the active lever but not of the inactive lever (two-way RM ANOVA, session × lever interaction, F_(8,64)_ = 7.417, p < 0.0001) (**Fig. 4b,d**). For the active lever, total presses were higher during each of sessions 2-9 than during session 1 (p = 0.0011 for session 2 vs. session 1; p < 0.001 sessions 3-9 vs. session 1, Dunnett’s multiple comparisons test). Conversely, inactive lever presses were not significantly different during sessions 2-9 compared with session 1. Mice injected with a control virus encoding EGFP did not acquire lever pressing over the course of eight training sessions (two-way RM ANOVA, session × lever interaction, F_(7, 21)_ = 1.484, p = 0.23) (**Fig. 4c**). Varying stimulation rates in optical self-stimulation experiments frequently results in modulation of lever-pressing behavior. To test the influence of stimulation rate on responding for thalamostriatal stimulation, we varied the stimulation train to either 20 pulses delivered at 20 Hz or 5 pulses delivered at 5 Hz (**Fig. S4d**) and performed within-subject comparisons of presses per session under baseline (10-Hz) and 20-Hz or 5-Hz stimulation conditions. Varying the stimulation train reduced the number of presses per session for both the 20-Hz (p = 0.226, paired t-test, **Fig. S4e**) and 5-Hz (p = 0.0148, **Fig. S4f**) conditions.

**Figure 4.**
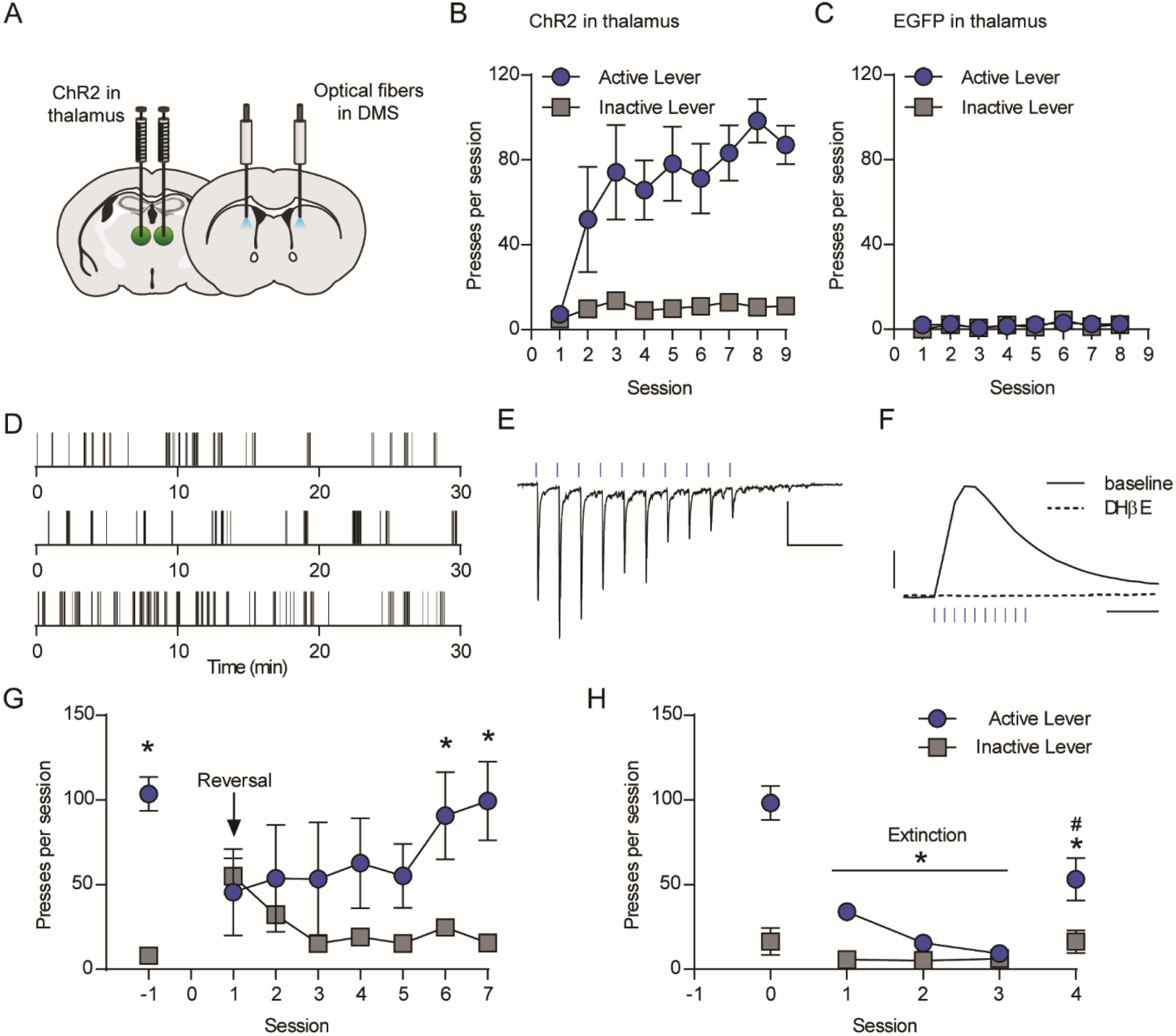
Optical stimulation of thalamic terminals in the DMS reinforces operant lever pressing. (a) Schematic of AAV-ChR2 injection in the anterior intralaminar nuclei of the thalamus and optical fiber placement in the DMS. (b-c) Acquisition of lever pressing for 10 pulses of 10-Hz optical stimulation of thalamostriatal terminals in the DMS of C57BL/6J mice expressing ChR2 (b, n = 8) or EGFP (c, n = 4). (d) Examples of lever presses from 3 representative mice expressing ChR2 in thalamostriatal terminals during acquisition session 9. Tick marks represent individual lever presses. (e) Representative whole-cell recording of EPSCs in a DMS medium spiny neuron in response to 10 pulses of 10-Hz optical stimulation. Scale bars: 100 pA, 0.25 s. (f) Representative FSCV traces of dopamine release measured in the DMS in response to 10 pulses of 10-Hz optical stimulation, before and after bath application of DHβE (1 µM). Scale bars: 0.5 µM dopamine, 0.5 s. (g) Lever presses at baseline and across 7 sessions of reversal learning (n = 5). *p < 0.05, active lever vs. inactive lever, Sidak’s multiple comparisons test. (h) Lever presses at baseline, during 3 extinction sessions, and during one reinstatement session (n = 5). *p < 0.05, active lever presses during extinction/reinstatement sessions vs. baseline, Tukey’s multiple comparisons test. For (b-c) and (g-h), data represent mean ± SEM.

Brain slice electrophysiology and FSCV experiments confirmed that optical stimulation in DMS stimulated both glutamate and dopamine release. In MSNs, the 10-Hz stimulation train reliably produced excitatory postsynaptic currents (**Fig. 4e**). In addition, 10-Hz stimulation of thalamostriatal terminals evoked dopamine release that was blocked by DHβE, which is consistent with dopamine release being driven by thalamic activation of CINs and subsequent acetylcholine actions on dopaminergic terminals (**Fig. 4f**) (Cover et al., 2019; Threlfell et al., 2012).

Next, we evaluated the ability of mice to flexibly update rates of lever pressing for thalamostriatal stimulation in response to changes in the lever-stimulation contingency. First, we reversed the lever-stimulation contingency such that the previously inactive lever became the active lever. Upon reversal of the active lever, mice decreased pressing of the formerly active lever and increased pressing of the newly active lever across seven training sessions (two-way RM ANOVA, lever × session interaction, F_(7,28)_ = 4.554, p = 0.0017) (**Fig. 4g**). By the sixth and seventh reversal training sessions, mice pressed the newly active lever more than the previously active lever (session 6, p = 0.0022; session 7, p = 0.0001, Sidak’s multiple comparisons test). We then evaluated responding under extinction conditions for three sessions followed by a single session in which we reinstated the lever-stimulation contingency. When stimulation was no longer delivered in response to a press on the previously active lever, mice rapidly decreased responding; restoration of press-stimulation contingency during a single reinstatement session restored pressing to 52.1 ± 7.5% of baseline levels (**Fig. 4h**). Across baseline, extinction, and reinstatement sessions, two-way RM ANOVA revealed a significant lever × session interaction (F_(4,16)_ = 23.87, p < 0.0001). *Post hoc* comparisons demonstrated that mice pressed the active lever less during each of the three extinction sessions as well as on the reinstatement day compared with baseline pressing (p < 0.0001, Tukey’s multiple comparisons test). When the lever-stimulation contingency was reinstated, mice pressed the active lever significantly more than during the final extinction session (p < 0.0001), but less than on the baseline day (p < 0.0001). Compared with the baseline session, there were no significant differences in responding on the inactive lever on any extinction day or during the reinstatement session

### Pharmacological manipulation of group II mGlu receptors modulates lever pressing for thalamostriatal stimulation

In the same group of mice trained to press a lever for thalamostriatal stimulation, we evaluated the ability of pharmacological manipulations of group II mGlu receptors to modulate lever pressing. Pharmacological interventions were performed after the acquisition and stimulation rate manipulations but prior to the reversal learning and extinction training shown (**Fig. S4a**). Injection of the group II mGlu receptor antagonist LY341495 (3 mg/kg) increased total lever presses to 209 ± 23.7% of vehicle pressing (p = 0.045, paired t-test) (**Fig. 5a, d-e**). Conversely, the agonist LY379268 dose-dependently reduced responding (1 mg/kg: 78.5 ± 10.2% of vehicle, p = 0.037; 3 mg/kg: 39.1 ± 10.7% of vehicle, p = 0.0023) (**Fig. 5b-d, f**). Detailed analysis of patterns of lever presses revealed that LY341495 and LY379268 had opposing effects on the number of clusters of pressing per session, with LY341495 increasing and LY379268 decreasing the number of clusters (paired t-test, LY341495: p = 0.0039; LY379268: p = 0.0133) (**Fig. 5g,l**). Similar to the effects of LY379268 on operant responding in Vglut2-Cre^+/−^;Ai32^+/−^ mice, the architecture of pressing within clusters was not consistently altered when considering cluster duration, presses per cluster, or within-cluster press rate (**Fig. 5m-o**). In addition, we did not observe a significant increase in break length (**Fig. 5p**). Within-cluster parameters and break length were similarly unaffected by LY341495 (**Fig. 5h-k**).

**Figure 5.**
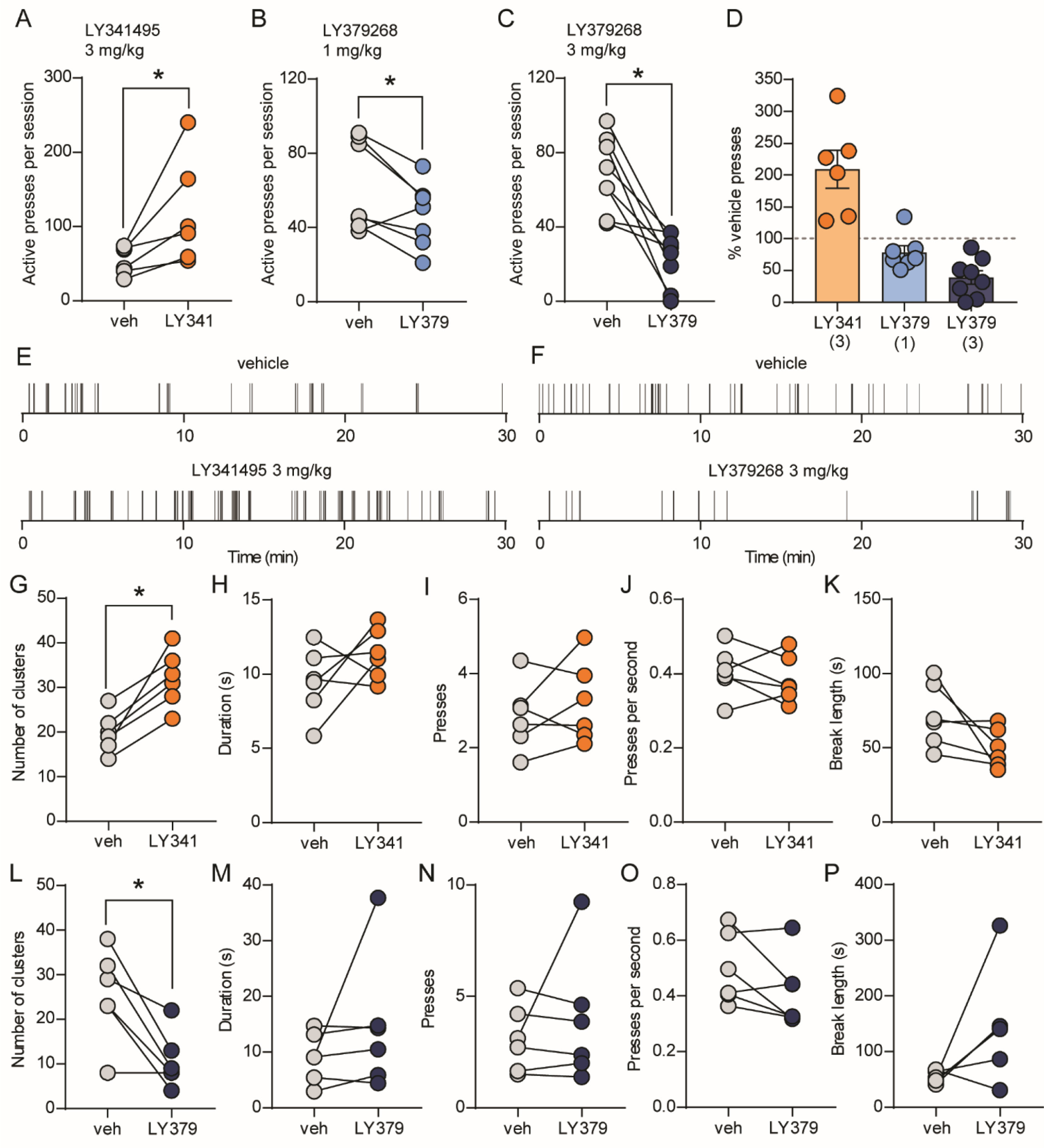
mGlu_2/3_ activity constrains operant responding for thalamostriatal stimulation. (a-c) Within-subject comparisons of lever presses per session in C57BL/6J mice expressing ChR2 in thalamostriatal terminals (n = 6-8) after injection of vehicle or 3 mg/kg LY341495 (a), 1 mg/kg LY379268 (b), and 3 mg/kg LY379268 (c). (d) Average active lever presses per session (normalized to vehicle presses for each mouse) for each drug treatment. Bars represent mean ± SEM, and individual data points are overlaid. (e-f) Examples of lever presses from a representative mouse during vehicle or drug sessions. Tick marks represent individual lever presses. (g-p) Within-subject comparisons of patterns of pressing during vehicle vs. 3 mg/kg LY341495 (g-k) or vehicle vs. 3 mg/kg LY379268 (l-p) sessions. Parameters analyzed were clusters of pressing per session (g,l), mean duration of clusters (h,m), mean number of presses per cluster (I,n), mean within-cluster press rate (j,o), and the mean length of breaks between clusters of pressing (k,p). *p < 0.05, paired t-test.

### Emergence of movements associated with optical stimulation

In both Vglut2-Cre^+/−^;Ai32^+/−^ mice and C57BL/6J mice injected with ChR2-encoding virus in the thalamus, we observed movements associated with optical stimulation of terminals in the DMS. Movements varied in type and magnitude both within and between animals, and included head turns, rearing, and partial or full rotations. Movements were not observed during passive delivery of stimulation trains during shaping or during the first session in which the mouse began lever pressing, but emerged by the third session in which the mouse pressed the lever, suggesting that the movements represent conditioned responses (**Supplemental video 1**). Interestingly, electrical stimulus-driven ICSS of dorsal striatum sites has been reported to induce similar movements (Phillips et al., 1976).

## DISCUSSION

Recent studies examining the roles of thalamostriatal projections in reinforcement learning have identified roles for discrete components of this pathway (i.e. inputs from the rostral intralaminar or parafascicular nuclei) in behavioral flexibility and incubation of drug craving (Bradfield et al., 2013; Kato et al., 2018; Li et al., 2018). In addition, optical stimulation of rostral intralaminar thalamic terminals in the dorsal striatum maintains operant responding in food-restricted mice previously trained to lever press for food (Cover et al., 2019). Here we demonstrate that specific stimulation of thalamic terminals in the DMS supports operant conditioning in a self-paced task. Mice acquire thalamostriatal self-stimulation in the absence of predictive cues, without manipulation of motivational state (food restriction), and without prior training to respond for an alternative outcome. Our findings extend upon previous reports that manipulations localized to the dorsal striatum, including stimulation of MSNs or nigrostriatal dopamine release, are sufficient to reinforce a self-initiated action (Ilango et al., 2014; Kravitz et al., 2012; Lalive et al., 2018; Rossi et al., 2013; Vicente et al., 2016).

In the nucleus accumbens, dopamine release caused by dopamine neuron spiking vs. local regulatory mechanisms is associated with discrete aspects of behavior, where local regulation of dopamine release tracks motivational state, while dopamine neuron spiking acts as a learning signal (Berke, 2018; Mohebi et al., 2019). To date, distinct roles for local regulation of dopamine release in the dorsal striatum have not been defined. Concordant with the idea that activation of intralaminar thalamus projections to the dorsal striatum produce behaviorally-relevant local regulation of dopamine release, inhibition of thalamostriatal neurons reduces presynaptic calcium signals in dopaminergic terminals while modestly decreasing spontaneous locomotor activity (Cover et al., 2019). In addition, projections from the rostral intralaminar nuclei to the DMS support methamphetamine seeking following forced abstinence in a D_1_ receptor-dependent manner (Li et al., 2018). The recent report by Cover et al. (2019) that D_1_ receptor antagonists decrease operant responding for thalamostriatal stimulation argues that locally-evoked dopamine release plays a critical role in reinforcement (Cover et al., 2019). This idea is supported by our finding that mGlu_2_-targeted pharmacological manipulations, which decrease thalamically-driven dopamine release, reduce operant responding for thalamostriatal stimulation. In Vglut2-Cre^+/−^;Ai32^+/−^ mice, we find that optical stimulation evokes dopamine release driven by CIN-mediated activation of nicotinic receptors on dopamine neurons as well as other mechanisms, most likely direct stimulation of dopaminergic terminals. Because dopamine release evoked by CIN activation constrains release driven by direct dopamine neuron activation (Threlfell et al., 2012), it is likely that in Vglut2-Cre^+/−^;Ai32^+/−^ mice, the glutamate-dependent, DHβE-sensitive component of dopamine release meaningfully influences dopamine release dynamics during self-stimulation. Importantly, our demonstration that mice specifically expressing ChR2 in thalamic inputs to the DMS also acquire self-stimulation behavior confirms that selective activation of glutamatergic inputs is sufficient to drive reinforcement.

Non-dopaminergic effects of glutamate released from thalamic terminals could also contribute to thalamostriatal self-stimulation. Notably, reinforcement learning mediated by optogenetic stimulation of DMS MSNs is independent of dopamine receptor activation (Kravitz et al., 2012). Thalamic inputs target both D_1_- and D_2_-expressing MSNs (Doig et al., 2010; Huerta-Ocampo et al., 2014) and drive excitation (Ellender et al., 2013; Lacey et al., 2007), raising the possibility that thalamostriatal self-stimulation is at least partially supported by direct activation of MSNs. Of note, ablation of CINs surrounding the site of thalamostriatal self-stimulation only partially impairs responding, suggesting involvement of mechanisms independent of locally evoked dopamine release (Cover et al., 2019). Direct optogenetic activation of D_1_-expressing MSNs in the DMS supports reinforcement learning (Kravitz et al., 2012; Lalive et al., 2018), whereas optical stimulation of D_2_-expressing MSNs promotes avoidance (Kravitz et al., 2012). Thus, in addition to local control of dopamine release, concurrent stimulation of excitatory inputs to both populations of MSNs could produce competing reinforcing and aversive signals that are reflected in the intermittent patterns of self-stimulation we observed.

Following from our previous report that mGlu_2_ activation reduces thalamostriatal glutamatergic transmission in both MSNs and CINs, and in turn reduces locally evoked dopamine release mediated by acetylcholine (Johnson et al., 2017), we predicted that pharmacological manipulation of mGlu_2_ would modify the reinforcing properties of light trains during our self-stimulation task. Supporting this, the mGlu_2/3_ agonist LY379268 reduced thalamostriatal self-stimulation in both Vglut2-Cre^+/−^;Ai32^+/−^ mice and C57BL/6J mice expressing ChR2 in the thalamus. In Vglut2-Cre^+/−^;Ai32^+/−^ mice, this effect was mimicked by the mGlu_2_-selective positive allosteric modulator BINA. Moreover, our finding that the mGlu_2/3_ antagonist LY341495 increased responding suggests that mGlu_2_ is endogenously activated during selective thalamostriatal terminal stimulation and constrains the reinforcing properties of stimulation. In contrast, we did not observe a significant increase in self-stimulation behavior in Vglut2-Cre^+/−^;Ai32^+/−^ mice following LY341495 administration, possibly owing to the different complement of terminals stimulated in this model vs. more specific expression of ChR2 in the intralaminar thalamic nuclei. However, an alternative explanation is that there is a smaller window to increase responding due to higher baseline rates of pressing observed in Vglut2-Cre^+/−^;Ai32^+/−^ mice.

Our finding that mGlu_2_ activation constrains operant responding for thalamostriatal stimulation identifies a unique neural substrate by which mGlu_2_ can modify the value of a stimulus during reinforcement learning. Activation of group II mGlu receptors is known to reduce both basal and psychostimulant-evoked dopamine release (Bauzo et al., 2009; D’Souza et al., 2011; Hu et al., 1999; Kim et al., 2005; Pehrson and Moghaddam, 2010). However, mGlu_2/3_ agonist administration does not reduce extracellular dopamine levels or locomotion evoked by midbrain electrical stimulation or L-DOPA administration (Pehrson and Moghaddam, 2010). These data are consistent with previous reports that failed to observe mGlu_2_ expression in nigrostriatal projections (Testa et al., 1998) and our current demonstration that LY379268-mediated inhibition of dopamine release is occluded by nicotinic receptor blockade in Vglut2-Cre^+/−^;Ai32^+/−^ mice. Collectively, these findings oppose the possibility that mGlu_2_ acts directly on dopamine neuron terminals to reduce dopamine release evoked by thalamostriatal stimulation or psychoactive drugs. In our thalamostriatal self-stimulation model, it is likely that mGlu_2_ acting on glutamatergic inputs to CINs, and possibly MSNs, underlies the dampened reinforcing properties of Vglut2^+^ or thalamostriatal terminal stimulation following administration of LY379268 or the mGlu_2_-selective PAM BINA. However, mGlu_2_ (and possibly mGlu_3_) actions in downstream circuit elements that support thalamostriatal self-stimulation could also contribute to these effects.

Consistent with the ability to decrease drug-enhanced extracellular dopamine levels, mGlu_2/3_ activation decreases self-administration of psychoactive drugs including cocaine, amphetamines, nicotine, and alcohol (Augier et al., 2016; Bauzo et al., 2009; Crawford et al., 2013; Dhanya et al., 2014; Dhanya et al., 2011; Jin et al., 2010; Justinova et al., 2016; Justinova et al., 2015; Kim et al., 2005; Li et al., 2016; Liechti et al., 2007; Sidhpura et al., 2010). However, the ability of these receptors to constrain reinforcement appears dependent on the nature of the reinforcer. In previous studies, mGlu_2/3_ agonists reduced responding for natural reinforcers such as sucrose, although such findings are inconsistent and typically occur at higher doses than are required to decrease responding for psychoactive drugs (Johnson and Lovinger, 2016; Justinova et al., 2016; Kufahl et al., 2011). Here we observed a modest decrease in responding for palatable food following LY379268 administration. Notably, administration of the mGlu_2/3_ antagonist LY341495 produced opposing effects on responding for thalamostriatal stimulation (increased responding) vs. food reinforcement (decreased responding). Similar to effects on thalamostriatal self-stimulation, previous studies have shown that LY341495 administration or genetic deletion of mGlu_2_ increases self-administration of reinforcing drugs such as alcohol, cocaine, and heroin (Gao et al., 2018; Morishima et al., 2005; Yang et al., 2017; Zhou et al., 2013). Inconsistent effects of mGlu_2/3_ manipulations on operant behaviors driven by different outcomes likely reflects differential engagement of circuitry modulated by mGlu receptors depending on the stimulus. Future studies employing techniques that permit observation of thalamostriatal activity and striatal dopamine dynamics during reinforcement learning will be necessary to determine the engagement of this pathway during acquisition of operant responding for various outcomes, including natural reinforcers and psychoactive drugs.

## METHODS

### Animals

Male and female mice were 5-8 weeks old at the time of surgery and 10-14 weeks old when behavioral experiments began. C57BL/6J mice were purchased from the Jackson Laboratory (strain 000664; Bar Harbor, ME) and arrived in the housing facility at least one week prior to surgery. To produce mice in which ChR2 was expressed under the control of the Vglut2 promoter, Vglut2-IRES-Cre^+/−^ mice (Jackson Laboratories stock no. 028863) were bred with Ai32^+/+^ mice (Jackson Laboratories stock no. 024109) to produce Vglut2-IRES-Cre^+/−^ and Vglut2-IRES-Cre^−/−^ mice hemizygous for the Ai32 allele. Animals were housed in the Fishers Lane Animal Care facility managed by the National Institute on Alcoholism and Alcohol Abuse (NIAAA). Studies were carried out in accordance with the National Institutes of Health guide for the care and use of laboratory animals and were approved by the NIAAA Animal Care and Use Committee. Mice were housed on 12 hour light/dark cycle on ventilated racks in a temperature- and humidity-controlled room with *ad libitum* access to food and water, except in the case of the food restriction procedures used in mice trained on operant responding for food reinforcement. All behavioral assessments were conducted during the light phase.

### Drugs and viral vectors

Adeno-associated virus serotype 1 (AAV1) encoding channelrhodopsin-2 (ChR2-H134R) and enhanced yellow fluorescent protein (EYFP) under control of a CamKII promoter was obtained from Addgene (plasmid #26969; Watertown, MA). Picrotoxin was obtained from Sigma-Aldrich (St. Louis, MO). Dihydro-β-erythroidine (DHβE) hydrobromide, LY341495, LY379268, and biphenyl-indanone A (BINA) were obtained from Tocris Bioscience (Minneapolis, MN). LY341495 (3 mg/kg) and LY379268 (1-3 mg/kg) were dissolved at 10 mM in 0.01N NaOH, diluted to dosing concentration in sterile water, and pH adjusted to ∼7. BINA (15 mg/kg) was dissolved in 10% Tween 80 and sonicated to form a microsuspension, then pH adjusted to ∼7. Drugs were administered intraperitoneally in a volume of 10 ml/kg.

### Stereotaxic viral vector injection and optical fiber implantation

Mice were anesthetized with isoflurane and placed into a stereotaxic frame (David Kopf Instruments). Craniotomy and durotomy were performed above each viral injection and fiber implantation site. C57BL/6J mice were injected bilaterally with 300 nL AAV-ChR2 at a rate of 60 nL/min using a Hamilton syringe with a 32-gauge needle. We targeted the anterior intralaminar nuclei of the thalamus using coordinates (relative to bregma) 1.3 posterior, 0.5 lateral, and 3.8 ventral from brain surface. For all mice used in self-stimulation experiments, tips of optical fibers (5 mm long, 205 µm core, 0.22 NA, with ceramic ferrule; Thorlabs) were implanted bilaterally in the DMS using coordinates (relative to bregma) anterior 0.8, 1.4 lateral, and 2.2 ventral from brain surface. Fibers were secured to the skull using Teets denture material (Co-oral-ite Dental Mfg., Diamond Springs, CA). Mice were allowed to recover at least 4 weeks prior to behavioral testing. After completion of behavioral testing, mice were transcardially perfused with phosphate-buffered saline (PBS) followed by 4% paraformaldehyde prepared in PBS. Brains were sectioned coronally at 50 µm and imaged on a Zeiss Axiozoom microscope to assess EYFP expression and confirm correct optical fiber placement.

### Behavioral apparatus

All experiments were conducted in wide mouse operant conditioning chambers (Med Associates, Fairfax, VT). Each chamber was equipped with two levers and a house light. Cue lights were present but were not used in the tasks. For experiments involving optical stimulation, the chamber had a modified top to allow mice to be tethered via a branched fiber and commutator to a patch cable (Doric Lenses, Quebec, Canada) connected to a 473-nm laser (Laserglow Technologies, Toronto, Candada). The laser power was adjusted to be 5-7 mW at the terminal of each branched fiber. For experiments involving food reinforcement, chambers contained a food hopper equipped with an infrared beam to detect head entries. Operant conditioning protocols were controlled through MED PC IV software (Med Associates). Optical stimulation trains were generated using a Master-8 (A.M.P.I, Jerusalem, Israel) controlling a shutter (Thorlabs, Newton, NJ).

### Optical self-stimulation task

Before each training session, mice were connected to a branched optical fiber and allowed to habituate to the tether for ∼30 minutes prior to placement in the operant conditioning chamber. Shortly after placement in the chamber, the session began with illumination of the house light and insertion of the lever(s), which remained in place for the duration of the session. In experiments with two levers, the lever assigned as active during initial training (right or left) was counterbalanced within groups. Optical stimulation (5-20 Hz, 1 s, 5 ms pulse width) in response to active lever presses was delivered on a fixed-ratio 1 (FR1) schedule, except that lever presses made during delivery of light stimulation were not reinforced by an additional stimulation train. Inactive lever presses did not produce any outcome. Sessions lasted 30 minutes with an unlimited number of reinforcements. The end of the session was signaled by extinguishing the house light and retracting the lever(s). To encourage exploration of the area immediately surrounding the active lever during early training, we performed a shaping procedure beginning 20 minutes into the first training session. During shaping, mice received an investigator-delivered stimulation train upon each approach of the lever (within ∼2 cm) or when the mouse orientated towards the lever when it was already in close proximity. Shaping continued for up to 1 (Vglut2-Cre;Ai32 mice) or 3 (C57BL/6J mice expressing ChR2 in the thalamus) additional training sessions, beginning 5 minutes into the session, and was terminated after the mouse pressed the active lever 3 times in a one minute window. For Vglut2-IRES-Cre^+/−^;Ai32^+/−^ mice, 3/13 acquired pressing without shaping, and the rest required shaping for part of session 2. Of the 8 C57BL/6J mice expressing ChR2 in the thalamus that acquired stable performance in the self-stimulation task, 4/8 mice underwent shaping until session 2, 2/8 underwent shaping until session 3, and 2/8 underwent shaping until session 4. To match procedures in control mice, Vglut2-IRES-Cre^−/−^;Ai32^+/−^ mice and C57BL/6J mice injected with control virus received shaping for the maximum amount of time shaping was performed for ChR2-expressing mice (through the 2^nd^ or 4^th^ session, respectively). All Vglut2-IRES-Cre^+/−^;Ai32^+/−^ mice acquired pressing behavior and were included in analyses. Several C57BL/6J mice injected with ChR2 in the thalamus were excluded from analysis for the following reasons: 1) mice pressed the active lever less than 20 times over the first four training sessions (n= 12); 2) mice acquired pressing, but not at sufficiently high or stable rates for meaningful analysis of effects of drug challenges (<30 presses/session on day 9, n =5); and 3) loss of implanted fiber during training (n = 1). Analysis of ChR2 expression in a subset of mice excluded for insufficient pressing revealed low levels of ChR2 expression in most cases.

After stable responding was established, reversal learning was assessed in a subset of mice by reversing the lever that produced light delivery. Learning of the new lever-stimulation contingency was assessed over 7 consecutive training sessions. Once stable pressing was observed following reversal learning, mice underwent extinction training in which an active lever press no longer produced a stimulation. Responding under extinction conditions was measured during 3 consecutive sessions, followed by a single reinstatement session in which the active lever-stimulation contingency was restored, without use of any priming or cues.

### Operant responding for food reinforcement

C57BL/6J mice were gradually food restricted to 85-90% of free-feeding body weight prior to training and maintained at ∼90% through the remainder of the experiment. The training protocol we chose was designed to produce high rates of pressing for food reinforcement. The beginning of each session was signaled by illumination of the house light and insertion of one lever (with the exception of the hopper training session, during which no lever was present). The first day of training consisted of non-contingent delivery of 20-mg grain-based food pellets (BioServ) into a hopper to acclimate the mice to the new food and retrieval from the hopper. Mice then received five training sessions (one per day) in which presses on a single lever were reinforced with delivery of a food pellet on an FR1 schedule. Once responding on an FR1 schedule was well-established, mice were trained on an FR5 schedule for 3 sessions, then an FR10 schedule for 3 sessions. All FR1 and FR5 training sessions lasted 60 minutes or were terminated early if the mouse received 30 reinforcements to avoid satiation. The first FR10 session lasted 60 minutes with no restriction on the number of reinforcer deliveries, and the final two FR10 sessions lasted 30 minutes with unlimited reinforcer deliveries. In 30 minute sessions on an FR10 schedule, mice typically did not exceed 30 reinforcements per session. Upon termination of each session, the house light was extinguished and the lever was retracted. Limited amounts of regular chow were provided after each day’s training session.

### Experimental timeline

Mice underwent one training session per day. Vglut2-Cre^+/−^;Ai32^+/−^ mice underwent 6 days of training prior to assessment of drug effects on operant responding. Experimental timelines for operant responding for food reinforcement and operant self-stimulation in the cohort of C57BL/6J mice expressing ChR2 in the thalamus are shown in Figs. **S3a** and **S4a**. After initial training, but prior to drug challenges, mice received habituation injections 30 minutes prior to training for at least 3 sessions. Between drug challenges or tests of different stimulation rates, mice underwent at least two, but typically three regular training sessions with vehicle injections, and response rates were monitored for stability between sessions. The order of drug testing was LY341495 (3 mg/kg), LY327968 (1 mg/kg), LY379268 (3 mg/kg) for all mice. We chose to administer escalating doses of agonist to minimize confounding effects of tolerance after repeated dosing, which has been reported with LY379268 (Liechti et al., 2007).

### Brain slice preparation, whole-cell patch clamp recording, and fast-scan cyclic voltammetry recording (FSCV)

Coronal brain slices (250 µm thick) were prepared using a vibratome (Leica Microsystems) as previously described (Johnson et al., 2017). Mice were anesthetized with isoflurane, decapitated, and brains were rapidly removed and submerged in ice-cold cutting solution containing (in mM): 30 NaCl, 4.5 KCl, 1 MgCl_2_, 26 NaHCO_3_, 1.2 NaH_2_PO_4_, 10 glucose, and 194 sucrose, continuously bubbled with 95% O_2_/5% CO_2_. Slices were placed in a 32°C holding chamber containing artificial cerebrospinal fluid (aCSF) containing (in mM): 124 NaCl, 4.5 KCl, 2 CaCl_2_, 1 MgCl_2_, 26 NaHCO_3_, 1.2 NaH_2_PO_4_, and 10 glucose, 305-310 mOsm, continuously bubbled with 95% O_2_/5% CO_2_. After 30-45 minutes at 32°C, slices were incubated at room temperature for at least 30 minutes prior to beginning experiments.

Whole-cell voltage-clamp recordings were conducted as previously described (Johnson et al., 2017). Hemisected slices containing DMS were submerged in and continuously perfused with 30-32°C aCSF at a rate of ∼1.5 mL/min. Recording pipettes (2.0-4.0 MΩ resistance in bath) were filled with Cs-based internal solution (295-300 mOsm) containing (in mM): 120 CsMeSO3, 5 NaCl, 10 TEA-Cl, 10 HEPES, 5 QX-314, 1.1 EGTA, 0.3 Na-GTP, and 4 Mg-ATP; pH was adjusted to 7.25 using CsOH. Slices were visualized on a Zeiss Axioskop 2 microscope. Cells for whole-cell recordings were visualized using a 40x/0.8 NA water-immersion objective. Recordings were performed using a Multiclamp 700A amplifier (Axon Instruments). MSNs were voltage-clamped at −60 mV throughout the experiment. Excitatory postsynaptic currents (EPSCs) were optically evoked by trains of 470-nm, 2-5-ms pulse-width field illumination delivered via a High-Power LED Source (Thor Labs). All recordings were filtered at 2 kHz and digitized at 10 kHz (Digidata 1322A, Axon Instruments). 50 μM picrotoxin was included in the aCSF to block fast GABAergic transmission. Acquisition was performed using Clamplex 10.3 (Molecular Devices).

FSCV recordings were performed as previously described (Johnson et al., 2017). A glass-encased cylindrical carbon fiber (Goodfellow, PA; 7-μm diameter, 100-130-μm exposed length) was placed into the DMS. Trains of optical stimulation (473 nm, 2 ms pulse-width, 10 mW output) were delivered by placing an optical fiber (105-μm core diameter, 0.22 NA, Thorlabs, NJ) in apposition to the brain slice. Extracellular dopamine release was monitored by applying a triangular input waveform from −0.4V to +1.2V and back to −0.4V (versus an Ag/AgCl reference electrode immersed in the bath solution) through the carbon fiber electrode. Cyclic voltammograms were collected at 10 Hz using a Chem-Clamp (Dagan Corporation) and DEMON Voltammetry and Analysis software (Yorgason et al., 2011). Stimulation trains were applied every 5 min, and drugs were bath applied to slices after stable baselines were established. Following recordings, carbon fiber electrodes were calibrated to known concentrations of dopamine, and evoked dopamine concentrations were calculated from recorded currents.

### Data analysis

Data visualization and statistical analyses were performed using GraphPad Prism. Lever pressing across training sessions was analyzed using two-way repeated measures (RM) analysis of variance (ANOVA). Session and lever were treated as within-subject factors, and genotype differences were analyzed as a between-group factor. Significant factor effects were analyzed by *post hoc* Dunnett’s, Sidak’s, or Tukey’s multiple comparisons tests depending on the comparisons being made. Within-subject effects of drugs or changes in stimulation rate were analyzed using a paired t-test. To provide a clear presentation of the magnitude of drug effects, we report the effects of each drug normalized to individual animal’s vehicle session then averaged across subjects, but statistical analyses were performed on the raw number of presses per session using within-subject comparisons. Analyses of pressing architecture (parameters: clusters per session, presses per cluster, cluster duration, within-cluster press rate, and break length) were performed using a custom analysis program written in Python (code available upon request). Clusters of pressing were defined as bouts of multiple lever presses with inter-press intervals <10.0 seconds. Drug effects on peak dopamine release in FSCV experiments were analyzed using a paired t-test or two-way RM ANOVA. When averaged data are shown, data points or bars represent mean ± SEM, and points on bar graphs represent individual subjects. For all statistical comparisons, alpha was set at 0.05.

## Acknowledgements and contributions

We thank all members of the Lovinger laboratory for helpful discussions and Guoxiang Luo for assistance with genotyping. We thank Dr. David Kupferschmidt for technical advice and comments on the manuscript. This work was supported by NIAAA Division of Intramural Clinical and Biological Research ZIA AA000416 (D.M.L.) and NIH grant K99 AA025403 (K.A.J.). K.A.J. and D.M.L. conceived the project and wrote the manuscript. Y.M. and L.V. performed and analyzed FSCV experiments. K.A.J. performed and analyzed all other experiments.

