## Supplemental Figures for "Operant self-stimulation of thalamic terminals in the dorsomedial striatum is constrained by metabotropic glutamate receptor 2"

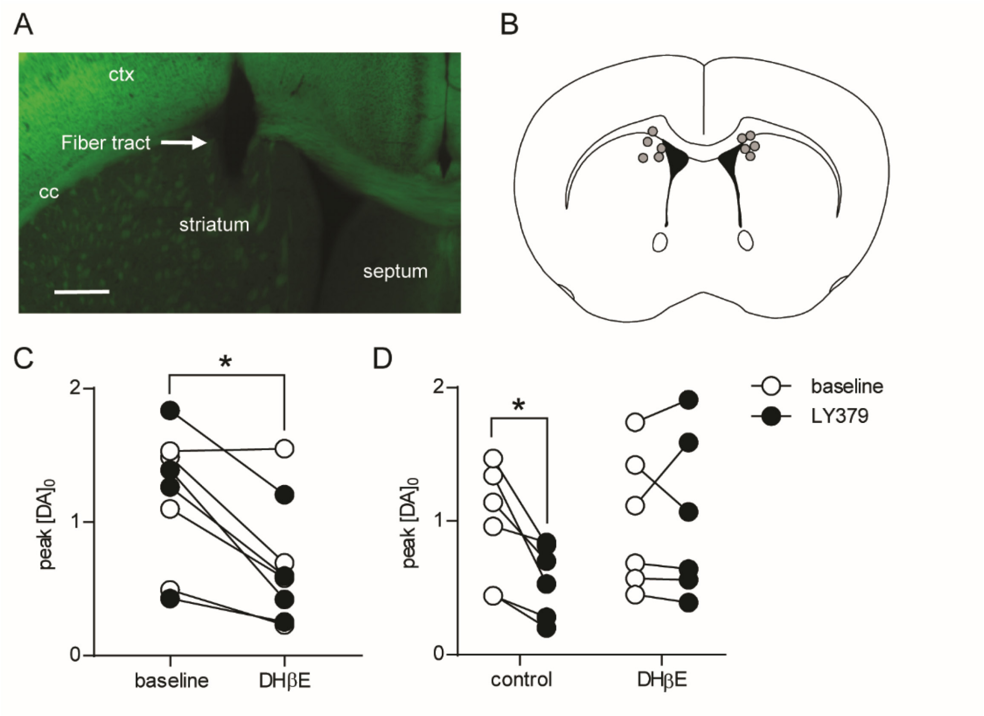


**Figure S1. Optical fiber placement and optically-evoked dopamine release in the DMS of Vglut2-Cre$^{+/-}$;Ai32$^{+}$ mice.** (a) Representative image of EYFP expression in a coronal section from a Vglut2-Cre$^{+/-}$;Ai32$^{+}$ mouse. Arrow indicates the optical fiber tract terminating in the DMS. Scale bar: 500 µm. (b) Schematic of optical fiber placement in mice represented in Figs. 1 and 2 (n = 5). (c) Peak dopamine concentrations before and after bath application of DHβE (1 µM) measured in response to 20 pulses of 20-Hz optical stimulation (n = 4, closed circles) or 10 pulses of 10-Hz stimulation (n = 4, open circles) in the DMS of Vglut2-Cre$^{+/-}$;Ai32$^{+}$ using FSCV. DHβE reduced peak dopamine responses to 56.2 ± 7.4% of baseline. *p = 0.0039, paired t-test. (d) Peak dopamine concentrations before (open circles) and after (closed circles) bath application of LY379268 (100 nM) in the absence or presence of DHβE (1 µM). Dopamine release was evoked by 10 pulses of 10-Hz stimulation. Two-way RM ANOVA revealed a main effect of LY379268 ($F_{(1,10)}$ = 5.518, p = 0.0407) and a significant LY379268 x DHβE interaction ($F_{(1,10)}$ = 7.363, p = 0.0218). *Post hoc* comparisons demonstrated that LY379268 reduced peak dopamine release in control slices, but not in the presence of DHβE (*p = 0.01 without DHβE; p = 0.96 with DHβE; Sidak's multiple comparisons test). Abbreviations: ctx, cortex; cc, corpus callosum.

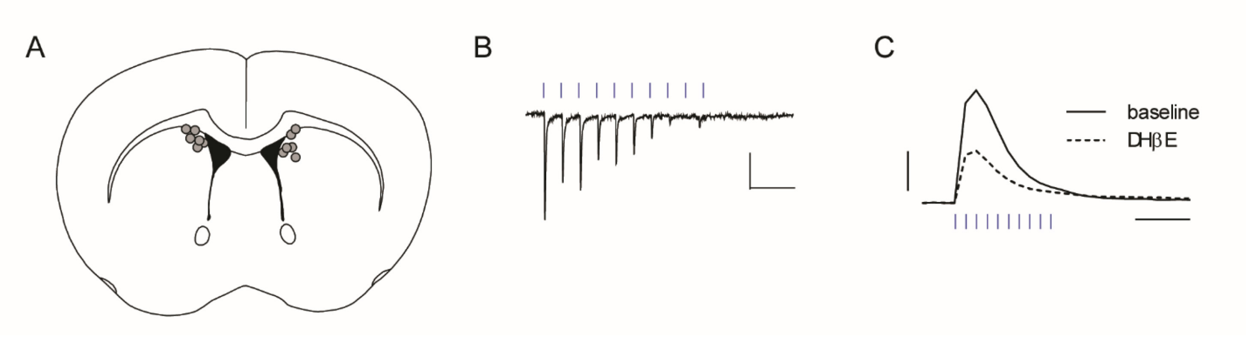


**Figure S2. Optical fiber placement and optically-evoked EPSCs and dopamine release in the DMS of Vglut2-Cre+/-;Ai32+ mice.** (a) Schematic of optical fiber placement in mice represented in Fig. 3 (n = 6). (b) Representative whole-cell recording of EPSCs in a DMS medium spiny neuron in response to 10 pulses of 10-Hz optical stimulation. Scale bars: 100 pA, 0.25 s. (c) Representative FSCV traces of dopamine release measured in the DMS in response to 10 pulses of 10-Hz optical stimulation, before and after bath application of DHβE (1 μM). Scale bars: 0.5 μM dopamine, 0.5 s.

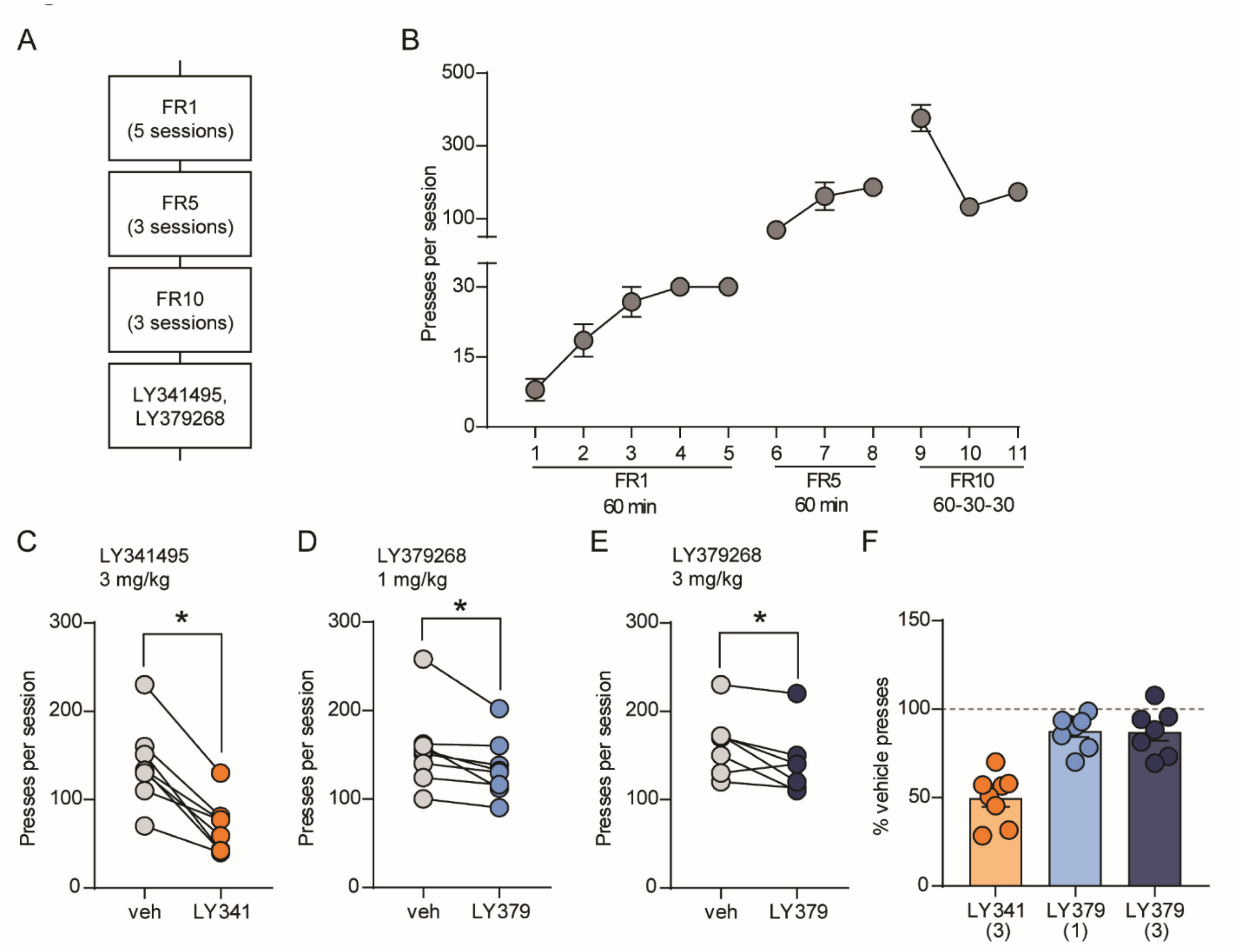


**Figure S3. Modulation of operant responding for food reinforcement by pharmacological manipulation of $mGlu_{2/3}$.** (a) Timeline of experimental procedures. (b) Acquisition of operant responding for food reinforcement over 11 training sessions (n = 8). (c-e) Within-subject comparisons of lever presses per session after injection of vehicle or 3 mg/kg LY341495 (c), 1 mg/kg LY379268 (d), and 3 mg/kg LY379268 (e) (n = 7-8). (f) Average lever presses per session (normalized to vehicle presses for each mouse) for each drug treatment. Bars represent mean ± SEM, and individual data points are overlaid. *$p < 0.05$, paired t-test.

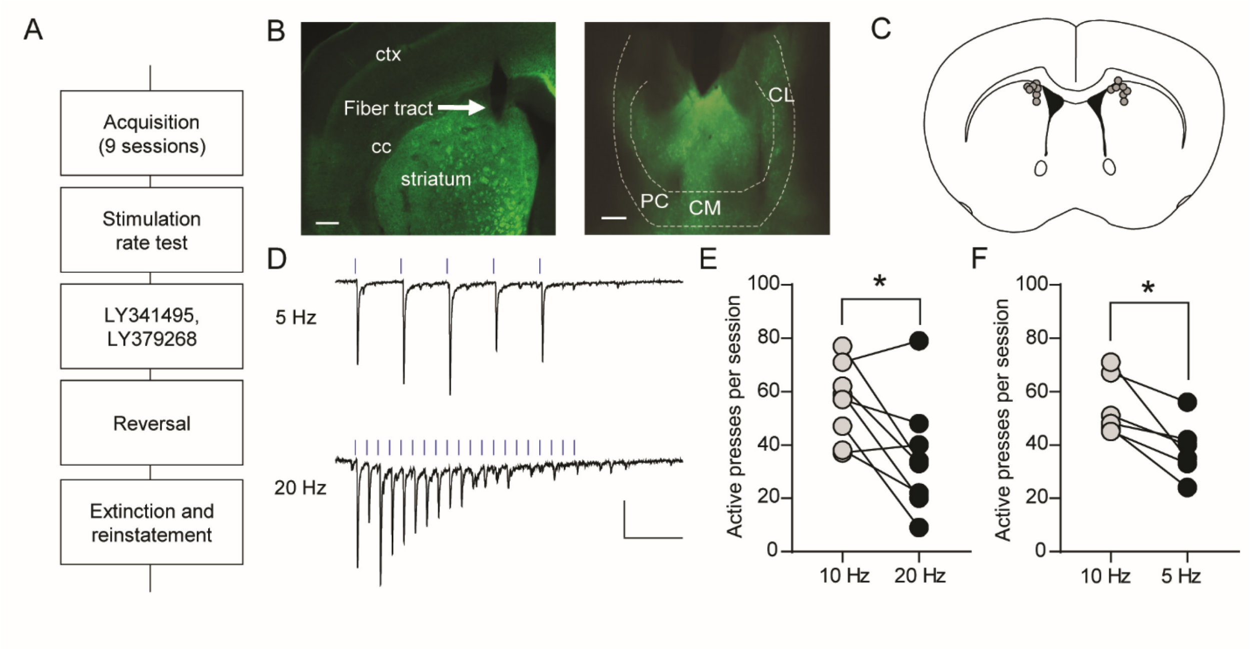


**Figure S4. ChR2 expression, optical fiber placement, and stimulation rate modulation of active lever pressing in C57BL/6J mice expressing ChR2 in thalamostriatal terminals.** (a) Timeline of experimental procedures. (b) Representative images of EYFP expression in the anterior intralaminar nuclei of the thalamus (right; scale bar: 250 μm) and optical fiber placement in the DMS (left; scale bar: 500 μm). ChR2 expression also extended posterior to the parafascicular nucleus in most animals. Arrow indicates the optical fiber tract terminating in the DMS. (c) Schematic of optical fiber placement in mice represented in Figs. 4 and 5 (n = 7). (d) Representative whole-cell recording of EPSCs in a DMS medium spiny neuron in response to 5 pulses of 5-Hz optical stimulation or 20 pulses of 20-Hz optical stimulation. Scale bars: 100 pA, 0.25 s. (e-f) Within-subject comparisons of active lever presses for 10-Hz vs. 20-Hz (e, n = 8) or 10-Hz vs. 5-Hz stimulation (f, n = 6). *$p < 0.05$, paired t-test. Abbreviations: ctx, cortex; cc, corpus callosum; CL, centrolateral nucleus of the thalamus; CM, centromedial nucleus of the thalamus; PC, paracentral nucleus of the thalamus.
